# Genetic analysis of bone morphometry and ivory vertebrae in threespine stickleback

**DOI:** 10.64898/2026.04.13.718284

**Authors:** Veronica C. Behrens, David Lee, Julia I. Wucherpfennig, David M. Kingsley

## Abstract

Previous genetic studies of skeletal variation in threespine stickleback fish (*Gasterosteus aculeatus*) have focused primarily on striking morphological differences. Here, we examine the largely unexplored genetic architecture of internal bone microstructural variation between marine and freshwater stickleback. µCT X-ray analysis revealed differences in the porosity, bone thickness, and bone volume fraction within armor plates and vertebrae from a marine and freshwater stickleback. Quantitative trait locus mapping in F2 progeny from a marine × freshwater stickleback cross identified a significant locus on chromosome 4 influencing multiple aspects of armor plate internal microstructure. This locus overlaps the well-characterized *Eda* region previously known to control armor plate number and size. Co-mapping of bone microstructure could be due to pleiotropic effects of *Eda* on multiple aspects of plate development or to changes in closely linked genes including *Itm2a*, which also plays a role in bone formation. Most bone microstructure traits in vertebrae showed weak or no genetic signal, consistent with a polygenic architecture. However, we identified a highly significant locus on chromosome 17 that is strongly associated with abnormally thickened “ivory vertebrae” that occurred in 8.4% of F2 offspring. This phenotype resembles Paget’s disease in humans, and the major locus region contains *Tnfrsf1b*, the stickleback ortholog of a human Paget’s disease susceptibility gene *TNFRSF11A*. Together, our findings identify genetic loci underlying natural variation in bone microstructure in wild fish and reveal a candidate gene associated with a disease-like skeletal phenotype, highlighting stickleback as a model for studying both evolutionary and pathological bone biology.

## Introduction

How and why certain traits evolve in natural populations is a longstanding question in evolutionary biology (Owen 1848; Darwin 1859; Endler 1986; Carroll 2005; Martin and Orgogozo 2013). The threespine stickleback, *Gasterosteus aculeatus*, is a uniquely powerful vertebrate model for crossing divergent populations and uncovering the genetic mechanisms underlying the evolution of skeletal traits. At the end of the last glacial period 10,000 to 20,000 years ago, stickleback from marine environments began colonizing newly-formed freshwater habitats. Confronted with similar ecological challenges, many different freshwater stickleback populations independently evolved similar traits (Bell and Foster 1994).

Despite dramatic skeletal differences, marine and freshwater stickleback retain the ability to interbreed. Genetic crosses can therefore be generated between ecotypes in the laboratory, producing first generation (F1) hybrids and second generation (F2) progeny that exhibit a range of phenotypes whose distribution can be compared to the inheritance of chromosome regions from the parents of the cross. Through quantitative trait locus (QTL) mapping using stickleback crosses, specific genetic loci have been identified that control striking morphological skeletal differences, including the complete loss or changes in size or number of dorsal spines (Howes *et al*. 2017; Wucherpfennig *et al*. 2022), lateral bony armor plates (Colosimo *et al*. 2004), and the pelvic girdle (Shapiro *et al*. 2004) in freshwater fish (Kingsley and Peichel 2006; Peichel and Marques 2017).

However, less attention has been given to bone internal microstructural differences (e.g. porosity of the bone, average size of the pores, average bone thickness) between marine and freshwater stickleback, in part because assessing these traits requires high resolution imaging techniques such as micro-computed tomography (µCT) or scanning electron microscopy (SEM). The few studies that have examined the microstructure of stickleback bones show that armor plates have a textured surface and internal structure containing heterogeneous pores (Song *et al*. 2010; Lees *et al*. 2012). Others have focused on population differences, demonstrating that the armor plates of freshwater stickleback are both more porous (Wiig *et al*. 2016) and have lower bone mineral density (Jamniczky *et al*. 2018) compared to marine populations. Armor plates are also major sites of phosphate accumulation in stickleback, and previous work has shown that genetic variation in armor plate development influences whole-body elemental ratios in wild populations (Durston and El-Sabaawi 2017).

In weight-bearing land animals including humans, increased porosity and decreased bone density cause the common disease osteoporosis, with an elevated risk of skeletal fractures (Compston *et al*. 2019). In contrast, evolved changes in skeletal porosity in aquatic animals may be an adaptive trait, helping to achieve neutral buoyant density in habitats with different salinities or altering body mass distribution, skeletal flexibility, and metabolic demands in environments with different predators and ion availabilities (Reimchen 1983; Taylor and McPhail 1986; Marchinko and Schluter 2007; Myhre and Klepaker 2009; Huang *et al*. 2024). Despite both the medical and evolutionary interest in bone density traits, the genetic basis of stickleback bone internal microstructural differences remains largely unexplored.

Here, we conduct QTL mapping using a cross between a marine and freshwater stickleback to investigate the genetic basis of internal bone morphometric variation assessed by µCT in both armor plates and vertebrae. By including armor plates, we build on the substantial body of literature focused on the evolutionary significance of lateral armor. Indeed, most previous bone morphometric studies in stickleback have focused on armor plates, and various hypotheses have been proposed for reduced bone number and size in freshwater stickleback that could also apply to bone internal microstructure variation. For example, reduced armor could provide an advantage in the face of grappling predation by macroinvertebrates (Reimchen 1992), lower availability of ions for bone formation in some environments (Giles 1983; Marchinko and Schluter 2007), or reduced buoyancy requirements in freshwater (Myhre and Klepaker 2009). We also include vertebrae in our study of internal bone microstructure to examine a skeletal region that is particularly relevant to changes in bone density and the clinical problems of prevalent osteoporosis and other bone disorders in humans (Kado *et al*. 2003; Rachner *et al*. 2011; Compston *et al*. 2019; Nguyen *et al*. 2024).

## Materials and methods

### Phenotyping using µCT scans

We initially µCT scanned wild-caught F0 stickleback fish to identify skeletal microstructure traits that differed between the marine and freshwater founders of a cross between a Bodega Bay (BDGB), California, USA, male and a Boulton Lake (BOUL), British Columbia, Canada, female (Fig. 1). We then scored the F2 progeny of the cross for a subset of the skeletal microstructure traits that varied between the F0 founder fish, as well as for other skeletal traits that could be easily measured by µCT.

**Figure 1.**
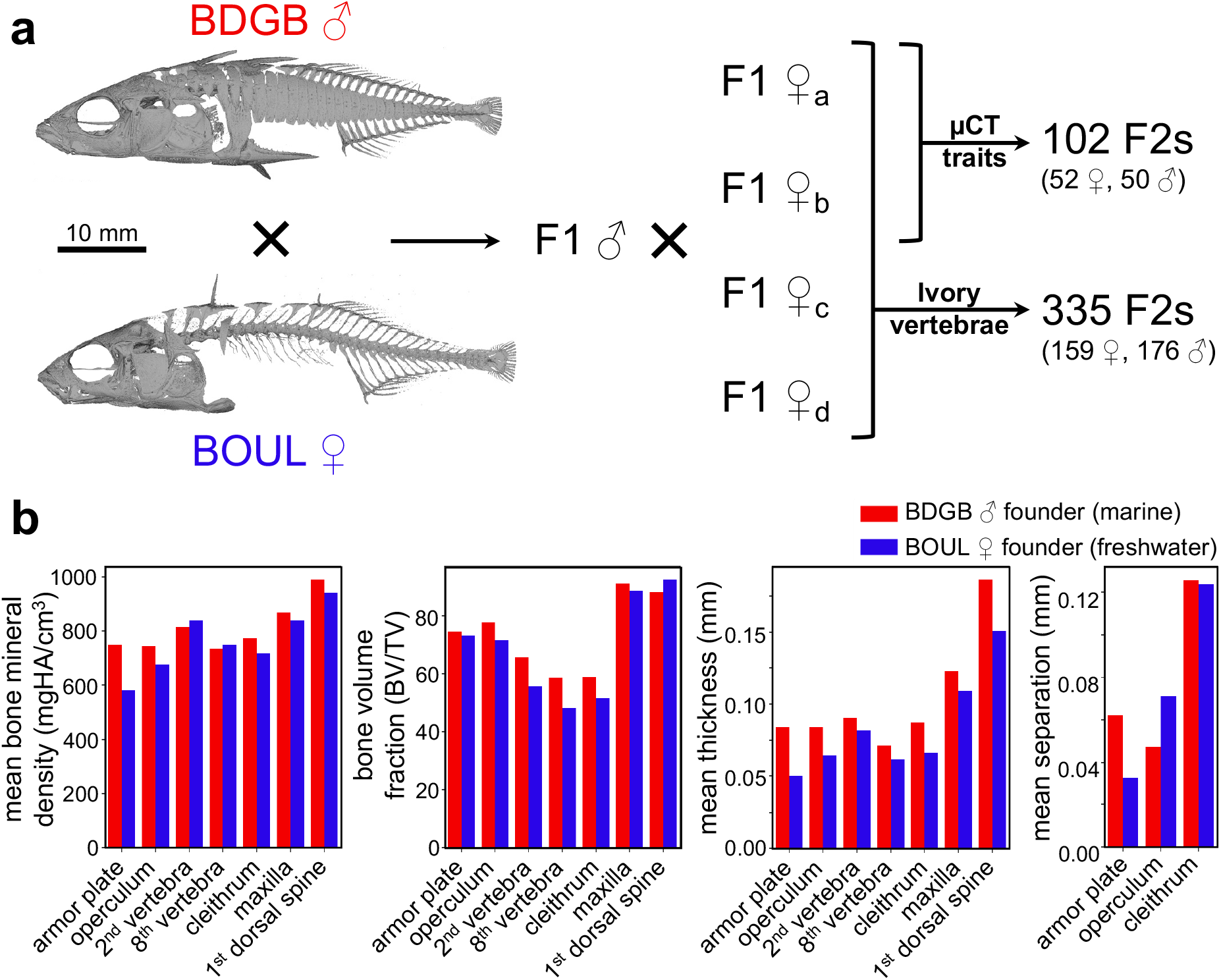
Genetic cross between a marine and freshwater stickleback with divergent bone morphometry traits. **a**.Genetic cross between a marine male (BDGB) and freshwater female (BOUL) F0 founder used for QTL mapping. A single F1 male was crossed to four F1 females. 102 F2s from two families were µCT scanned and scored for all traits. These same 102 F2s and 233 additional F2s from two other families were X-rayed and scored for ivory vertebrae. **b**.Quantification of bone morphometry traits for the F0 BDGB and BOUL founders. The two F0 founders were µCT scanned, voxels from each of 7 bones (x-axis) were isolated manually using Dragonfly imaging software, and bone morphometry measurements (y-axis) were calculated using Dragonfly’s Bone Analysis Extension. For most bones measured, the BOUL freshwater founder has lower mean bone mineral density, bone volume fraction (BV/TV), and mean bone thickness. Note that mean bone separation was calculated only for bones with obvious porous honeycomb-like internal structures.

CT scans were generated using a MicroCT42 (Scanco Medical AG) with the following parameters: 10 µm^3^ voxel size, 70 kVp, 114 µA, 8 W, 300 ms integration time, and frame averaging of 2. Whole stickleback were scanned in acrylic cylinders (20 mm diameter ×75 mm high) provided by the manufacturer filled with 70% ethanol and stabilized with tissue sponges (Epredia™ 8453).

For measuring bone morphometry traits, the raw scan was converted to a stack of DICOM images using the µCT Evaluation Program preinstalled on the workstation provided by the manufacturer and then transferred to an external workstation. The substack of DICOM images containing the bone-of-interest was loaded into Dragonfly version 2020.1 for Windows (Comet Technologies Canada Inc., Montreal, Canada; software available at https://dragonfly.comet.tech/) and further cropped using the clipping box. Then, Dragonfly’s Image Processing Panel was used to apply the slope map filter (scale = 10), mean shift algorithm, and isodata thresholding to segment bone from background. Voxels not belonging to the bone-of-interest were manually removed using a combination of Dragonfly’s 3D lasso tool, 3D painter tool, and 2D painter tool. Finally, Dragonfly’s Bone Analysis Extension was used to compute bone morphometry measurements in three steps.

#### Step 1

Holes in the original unfilled bone were filled (hole size of 0.3 mm or 0.1 mm for the armor plate and 2nd vertebra, respectively) followed by manual filling of larger holes missed by the algorithm.

#### Step 2

This step normally separates cortical bone from trabecular bone, but it was skipped completely in this analysis because it is unreliable for fish bones.

#### Step 3

The following measurements were computed and recorded for the bone-of-interest using 0.1 mm as the estimated mean bone thickness (Bouxsein *et al*. 2010) (Supplementary Fig. 1):

- Surface area: Surface area of the unfilled bone
- Total volume (TV): Volume of the filled bone
- Bone volume (BV): Volume of the unfilled bone
- Bone volume fraction (BV/TV): Ratio of the volume of the unfilled bone to the filled bone
- Mean bone thickness: Mean thickness of the unfilled bone, assessed in 3D. Note this is described as “Trabecular thickness” by Dragonfly’s Bone Analysis Extension.
- Mean bone separation: Mean distance between the honeycomb-like structures in the unfilled bone, assessed in 3D. This was assessed only for bones with obvious honeycomb-like internal structures including the armor plate, operculum, and cleithrum. Note this is described as “Trabecular separation” by Dragonfly’s Bone Analysis Extension.

All bone morphometry measurements except bone volume fraction (BV/TV) were standardized by multiple regression against standard length (measured by standard X-ray using Fiji version 2.0.088 (Schindelin *et al*. 2012)) and sex.

For scoring right pleural rib number, right epipleural rib number, supraorbital grooves, and frontal bone sculpture, the raw scan was segmented using the µCT Evaluation Program preinstalled on the workstation provided by the manufacturer to keep only voxels with bone mineral density (BMD) of at least 150 mgHA/mm^3^ (Gauss sigma = 0.8 and Gauss Support = 1). Using Image Processing Language (IPL), the resulting _SEG.AIM file was downsampled by 2X, re-thresholded, and converted to a small mesh model using the scale, threshold, and stl commands, respectively. The resulting STL file was transferred to an external workstation and viewed in 3D Slicer version 4.10.2 (Fedorov *et al*. 2012) as a 3D model. Supraorbital grooves and frontal bone sculpture were scored on a scale from 1-5, with 5 indicating the greatest prominence (Supplementary Fig. 2).

In cases where BMD is reported, raw attenuation values were converted to hydroxyapatite (HA) density using the following linear conversion determined by scanning a density phantom with rods of known HA density (0, 100, 200, 400, and 800 mgHA/mm^3^) provided by the manufacturer using the same scan parameters: *BMD* = [*raw*_*attenuation*_*value×* (*slope*/4096)] + *intercept*, where *slope* = 411.3990 and *intercept* =− 227.3230.

µCT phenotyping was performed by the same person to ensure consistency, and was done blinded to genotype of the F2 offspring.

### Phenotyping using standard X-rays

Armor plate number on the left side, standard length, and ivory vertebrae presence or absence were scored using standard X-rays taken with an UltraFocus X-ray cabinet (Faxitron) using the following parameters: 38 kV and 4.8 s. Standard length was measured using Fiji version 2.0.088 (Schindelin *et al*. 2012). Ivory vertebrae were scored as present if any vertebrae appeared abnormally bright white and/or deformed. X-ray phenotyping was performed by the same person to ensure consistency, and was done blinded to genotype of the F2 offspring.

### Sexual dimorphism analysis

For each trait, sexual dimorphism was assessed using Welch’s t-test to compare the trait values of all male F2s versus all female F2s after correcting for standard length. P-values were adjusted for multiple testing using the Bonferroni correction (m = 17 tests).

### QTL mapping

The marine male Bodega Bay, California, USA (BDGB) × freshwater female Boulton Lake, British Columbia, Canada (BOUL) genetic cross (Fig. 1a) used here—including the F2 fish, genotype information, and linkage map—was previously described by Wucherpfennig *et al*. 2022. Briefly, a wild-caught BDGB and a wild-caught BOUL stickleback were crossed by *in vitro* fertilization, and F1s were raised in the laboratory. Because a sufficiently powered mapping cross required more F2 progeny than a single female fish could produce, sperm from a single F1 male was cryopreserved and used to fertilize eggs from four F1 females, generating four F2 families with a shared paternal parent. F2s were raised in the laboratory for 1 year, euthanized, and preserved in 70% ethanol. DNA was extracted from fin clips taken from the F0 founders and F2s and genotyped using an Illumina GoldenGate genotyping array. Intensity data were processed using GenomeStudiov.2011 (Illumina), and uninformative or low-quality SNPs were removed. The linkage map was constructed with tmap version 0.686 (Tennekes 2018) using 343 F2s and 452 markers.

QTL analysis was performed with the scanone function using Haley-Knott regression in R/qtl version 4.2.2 (Broman *et al*. 2003) with sex and family (i.e. maternal F1 lineage) as covariates. A normal model was used for all traits except ivory vertebrae which used a binary model. For each trait, N = 10,000 permutation tests were performed to determine the logarithm of the odds (LOD) significance threshold (*a* = 0.05). With the exception of the ivory vertebrae trait, which could be rapidly scored from standard X-ray images and was therefore analyzed in 335 F2s, all other traits were measured in 102 F2s due to the substantially greater time required for µCT imaging, processing, and quantification. Percent of phenotypic variance explained (PVE) by each significant QTL was calculated using the following equation where N is the number of F2s: 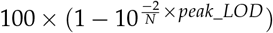.

### Identifying bone-related genes near peak QTLs

All stickleback genes in the regions surrounding peak QTLs and their corresponding human orthologs (Supplementary Fig. 4) were identified using Ensembl’s BioMart web tool (Smedley *et al*. 2009). The final set of bone-related genes was then determined by intersecting these human orthologs with the set of human genes annotated with one or more of the following Gene Ontology terms in the AmiGO online database (Ashburner *et al*. 2000; Carbon *et al*. 2009; The Gene Ontology Consortium *et al*. 2023): “ossification,” “bone development,” and “bone remodeling.”

### TNFRSF1B protein alignments and analysis

Human and medaka orthologs for stickleback *Tnfrsf1b* and their corresponding protein sequences and transmembrane helix domains were identified and downloaded using Ensembl’s BioMart web tool (Smedley *et al*. 2009). BDGB- and BOUL-specific amino acid changes in TNFRSF1B were identified using population-specific SNPs previously published in Roberts Kingman *et al*. 2021b originally discovered by DNA-sequencing 227 stickleback fish from 132 different populations. Protein alignments were performed using Geneious Prime® 2022.1.1 (https://www.geneious.com) via global alignment with free end gaps with the Blosum62 cost matrix. The SIFT web server (Sim *et al*. 2012) was used to predict the effect of amino acid changes on protein function.

## Results

### Bone morphometry differences between marine and freshwater stickleback

To study the genetic basis of stickleback internal bone morphometry differences, we used an F2 cross between a wild-caught BDGB marine male stickleback and a wild-caught BOUL freshwater female stickleback (Fig. 1a). This F2 cross was originally generated and genotyped to investigate the genetic basis of dorsal spine number and length variation (Wucherpfennig *et al*. 2022). We examined the BDGB and BOUL founders of the cross using three-dimensional high-resolution µCT scans and found that the F0 founder fish also differed in several internal bone morphometric traits (Fig. 1b). For the two founder fish, we manually isolated voxels from individual bones of several different types, including: flat porous bones (the armor plate between the first and second pterygiophore and the operculum); individual vertebrae in the spine (the 2nd and 8th); a bone with both a flat porous section and a long dense portion (the cleithrum); and long dense bones (the maxilla and the 1st dorsal spine). The most dramatic differences in bone mineral density, mean bone thickness, and mean bone separation were in the armor plate while the most dramatic difference in bone volume fraction (BV/TV) was in the vertebrae (Fig. 1b).

### Quantitative trait locus mapping of bone traits scorable by µCT

To examine the possible genetic basis of these internal bone microstructural differences, we also measured bone morphometric traits in armor plates and vertebra of 102 F2 progeny from the cross between BDGB and BOUL (Fig. 1a, Fig. 2a). We focused on the armor plate between the first and second pterygiophore (referred to simply as “the armor plate” for the remainder of this study) and on the 2nd vertebra. We isolated each bone from µCT scans and measured the following: surface area, total volume (TV), bone volume (BV), bone volume fraction (BV/TV), and mean bone thickness (Fig. 2a, Supplementary Fig. 1). Mean bone separation—the mean diameter of the pores within the bone—was calculated for the armor plate only. With the exception of bone volume fraction (BV/TV), these traits were standardized by multiple regression against standard length and sex.

**Figure 2.**
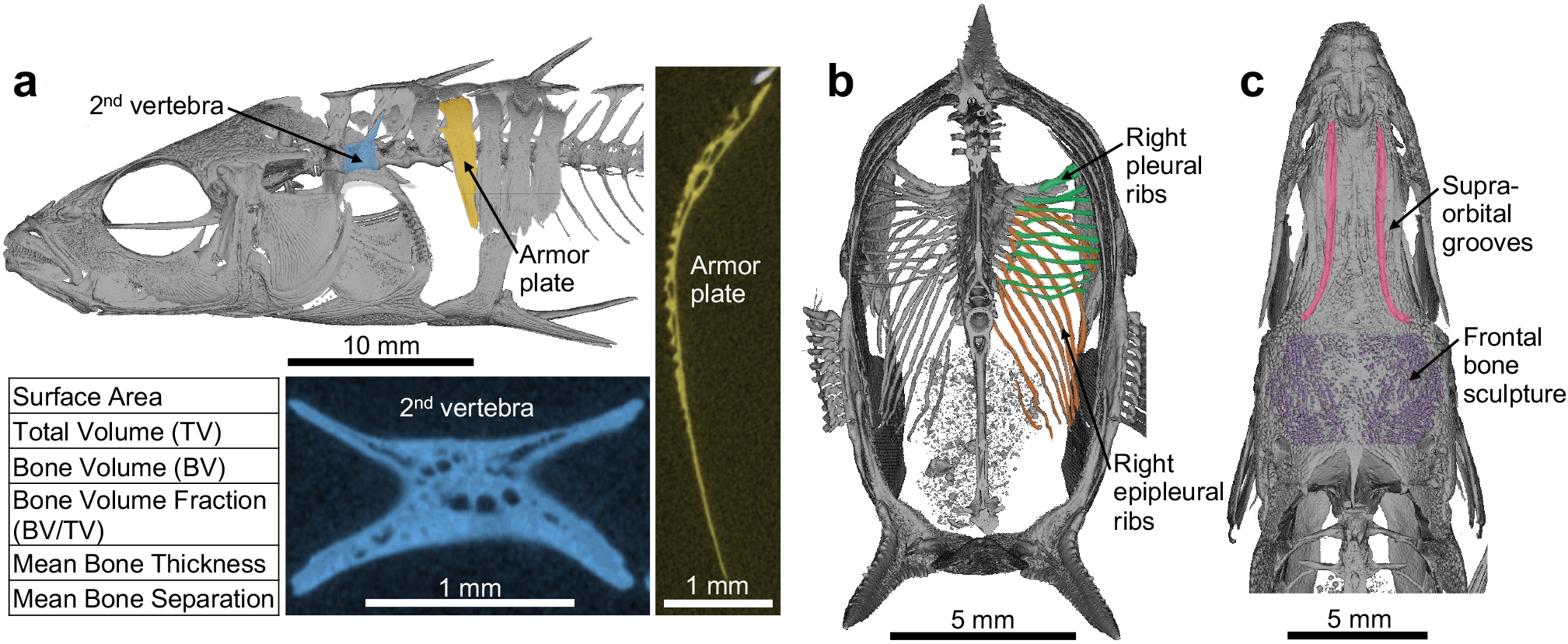
Bone morphometry traits scored via µCT for QTL mapping. **a**.(Top left) Lateral view of a representative F2 stickleback fish generated via µCT. The armor plate immediately posterior to the 1st dorsal spine is pseudo-colored in yellow, and the 2nd vertebra is pseudo-colored in blue. (Right) Representative µCT transverse cross section of the armor plate. (Bottom middle) Representative µCT longitudinal cross section of the 2nd vertebra. (Bottom left) Morphometry measurements scored for the armor plate and the 2nd vertebra for QTL mapping. All but bone volume fraction (BV/TV) were standardized by multiple regression against standard length and sex. Mean bone separation was scored for the armor plate only since the 2nd vertebra lacks obvious porous honeycomb-like internal structures. See Supplementary Fig. 1 for a schematic representation of µCT-derived bone morphometry measurements. **b**.Posterior view of a transversally-bisected representative F2 stickleback fish generated via µCT. Right pleural rib number (pseudocolored in green) and right epipleural rib number (pseudo-colored in orange) were counted for QTL mapping. **c**.Dorsal view of the skull of a representative F2 stickleback fish generated via µCT. The prominence of the supraorbital grooves (pseudo-colored in pink) and frontal bone sculpture (pseudo-colored in purple) were scored for QTL mapping on a scale from 1-5, with 5 indicating the greatest prominence. See Supplementary Fig. 2 for representative F2s from each semi-quantitative phenotype category.

We also scored several other traits that were easily measurable with µCT scans, including the total number of armor plates, the number of pleural ribs on the right side, the number of epipleural ribs on the right side, the prominence of the grooves on the dorsal side of the frontal bone (referred to as “supraorbital grooves”), and the prominence of texture on the dorsal side of the frontal bone (referred to as “frontal bone sculpture”) (Fig. 2b-c). Numerical traits were quantified by direct counting, and the two frontal bone traits were scored on a semiquantitative scale from 1-5, with 5 indicating the greatest prominence (Supplementary Fig. 2).

Multiple traits exhibited transgressive segregation, with some F2 fish displaying phenotype values beyond the range observed in either F0 founder fish, most notably among the vertebral traits (Fig. 3). This pattern is common in crosses between phenotypically divergent populations and can arise when recombination brings together complementary alleles or alleles with additive effects in novel combinations (Rieseberg *et al*. 1999). Differences in rearing environment could also have contributed to the more extreme F2 phenotypes, since the F0 founders were wild-caught while the F2 progeny were raised under laboratory conditions.

**Figure 3.**
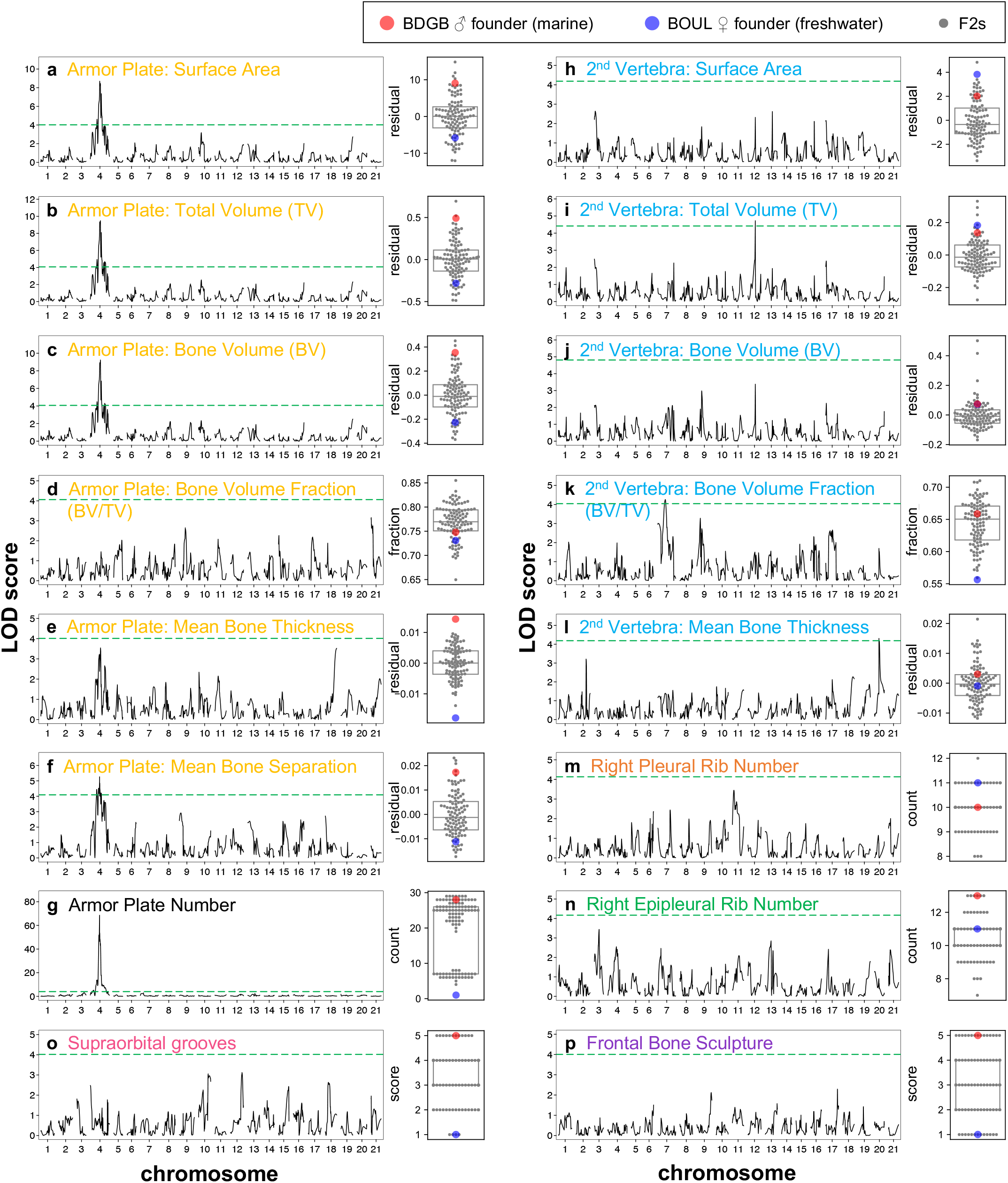
QTL mapping results for bone morphometry and other traits scored by µCT. Each panel (**a-p**) shows the QTL scan results (left) and a box plot of the distribution of phenotypes (right) for a single trait, with the title color corresponding to the color scheme in Fig. 2. The QTL scan results for each trait are represented by the logarithm of the odds (LOD) on the y-axis plotted against the chromosomal map position in cM on the x-axis, with the horizontal dashed green line indicating the genome-wide significance threshold (*α* = 0.05) calculated from 10,000 permutation tests. Each box plot represents the distribution of phenotypes observed for that trait, with each gray dot corresponding to the phenotypic value for a single F2 fish and the red and blue dot corresponding to the phenotypic value for the BDGB and BOUL F0 founder, respectively. Armor plate surface area (**a**), total volume (**b**), bone volume (**c**), and mean bone separation (**f**) map to the same region on chromosome 4 as armor plate number (**g**). 2nd vertebra total volume (**i**), bone volume fraction (**k**), and mean bone thickness (**l**) have QTLs that just pass the genome-wide significance threshold on chromosomes 12, 7, and 20, respectively. All other traits do not have significant detected QTLs. See Supplementary Fig. 3 and 4 for single-chromosome LOD plots of traits with genome-wide significant QTLs.

Using a linkage map of 451 markers (Wucherpfennig *et al*. 2022), we carried out QTL mapping to examine the genetic basis of these bone morphometry, bone number, and frontal bone traits scorable by µCT. We found a strong QTL on chromosome 4 for the following armor plate traits: surface area (LOD: 8.7, PVE: 32.5%), total volume (LOD: 9.5, PVE: 34.9%), bone volume (LOD: 9.2, PVE: 34.0%), and mean bone separation (LOD: 5.3, PVE: 21.3%) (Fig. 3a-c,f, Table 1, Supplementary Fig. 3). Armor plate number has been previously mapped to this same region of chromosome 4 (Colosimo *et al*. 2004), and this result is replicated for the 102 F2s used in this study (LOD: 68.6, PVE: 95.5%) (Fig. 3g, Table 1, Supplementary Fig. 3). We found only modest QTL peaks for 2nd vertebra traits, with the total volume (LOD: 4.7, PVE: 19.1%), bone volume fraction (LOD: 4.3, PVE: 17.6%), and mean bone thickness (LOD: 4.3, PVE: 17.6%) peaks just passing the significance threshold on chromosomes 12, 7, and 20, respectively (Fig. 3i,k-l Table 1, Supplementary Fig. 4). No significant QTL peaks were identified for any other traits, including rib number and frontal bone traits.

**Table 1.**
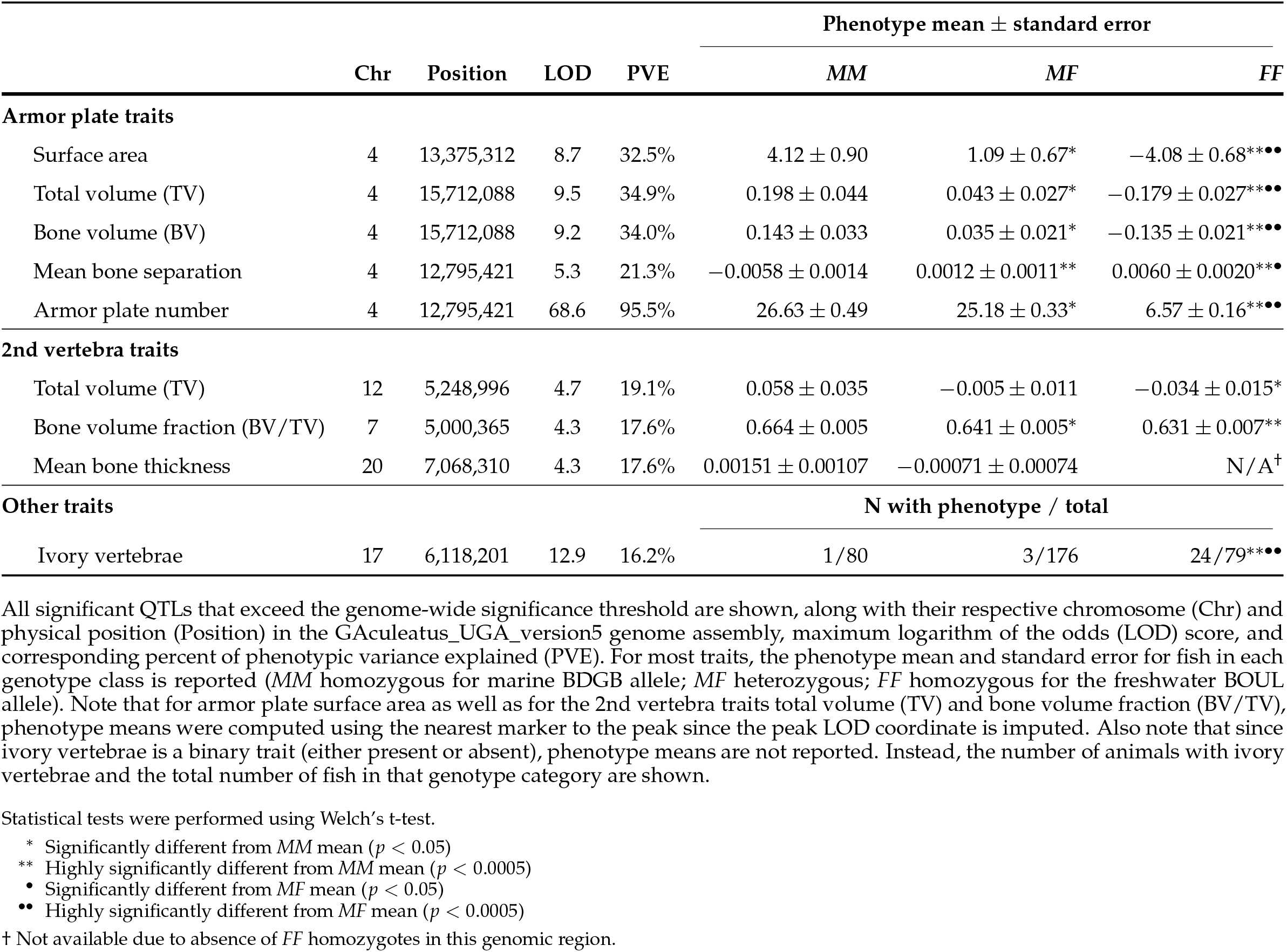
Significant QTLs.

### QTL mapping of ivory vertebrae

While analyzing standard two-dimensional X-rays from the same BDGB × BOUL cross, we noted that 28 of 335 F2s have abnormally bright white vertebrae (Fig. 4a). Similar bright white “ivory vertebrae” are also found in X-rays of humans with Paget’s disease, the second most common metabolic bone disease after osteoporosis (Kravets 2018). In the stickleback cross, both F0 founders were unaffected, and the number, location, and brightness of ivory vertebrae were highly variable across F2 progeny (Fig. 4a). µCT transverse cross-sections of the most commonly affected vertebra (vertebra 15, N = 7) showed overgrowth and abnormal morphology of the ivory vertebra compared to WT (Fig. 4b) and a statistically significant increase in mean bone mineral density (mean 829 mgHA/cm^3^ compared to 750 mgHA/cm^3^, *p* = 0.003 by Welch’s t-test) (Fig. 4c).

**Figure 4.**
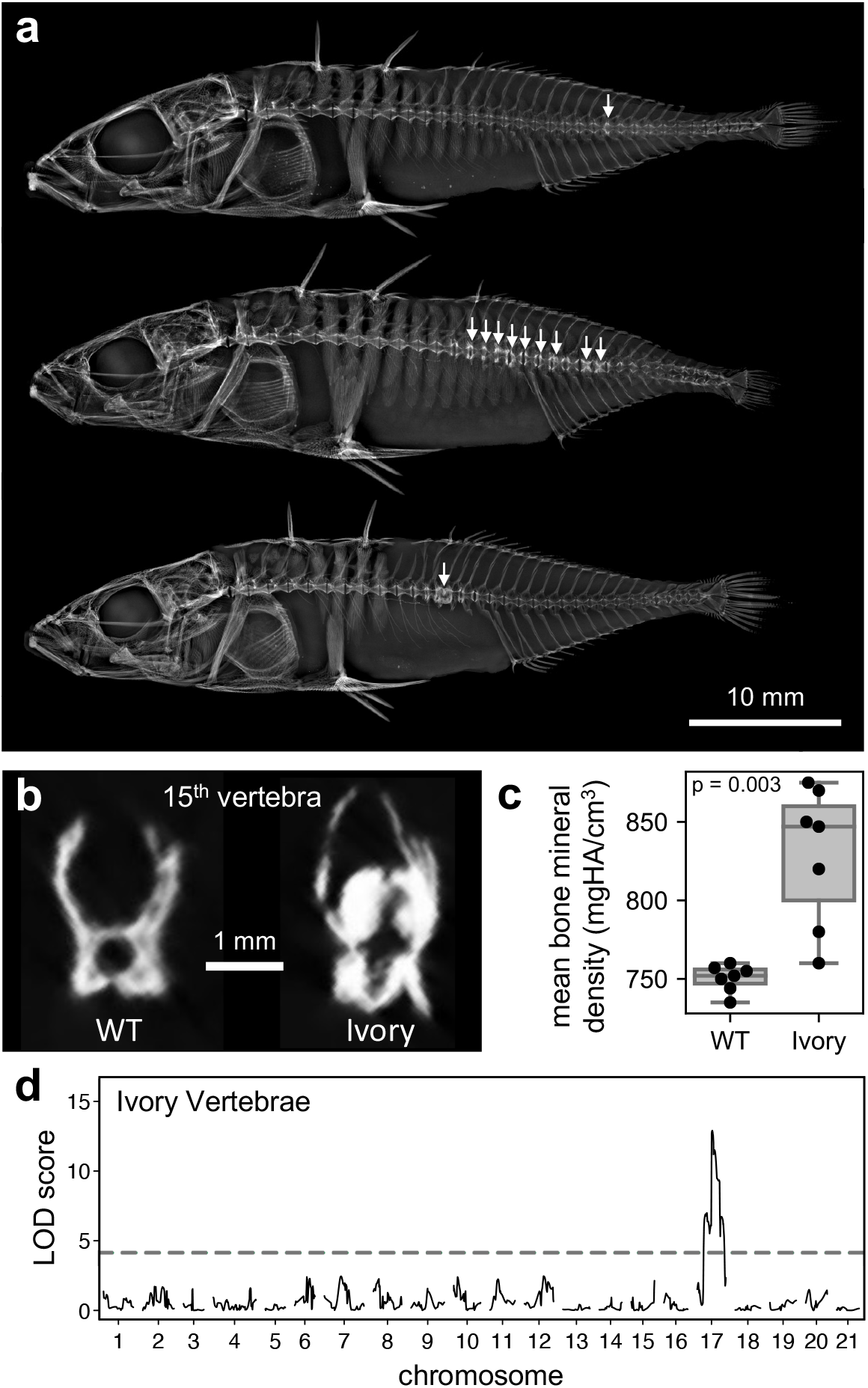
A major-effect QTL for ivory vertebrae on chromosome 17. **a**.Standard X-rays of representative F2 fish with white arrows indicating ivory vertebrae. **b**.Representative µCT transverse cross-sections of a wildtype (WT) and an ivory 15th vertebra. **c**.Quantification of mean bone mineral density (y-axis) for fish with a WT 15th vertebra (N = 7) and fish with an ivory 15th vertebra (N = 7). Ivory vertebrae have statistically significantly higher bone mineral density with a mean of 829 mgHA/cm^3^ compared to 750 mgHA/cm^3^ for WT vertebrae (*p* = 0.003 by Welch’s t-test). **d**.QTL scan results for ivory vertebrae showing a major-effect QTL on chromosome 17. Logarithm of the odds (LOD) is on the y-axis plotted against the chromosomal map position in cM on the x-axis, with the horizontal dashed line indicating the genome-wide significance threshold (*a* = 0.05) calculated from 10,000 permutation tests.

We identified a major-effect QTL for ivory vertebra presence/absence on chromosome 17 (LOD: 12.9) that explains 16.2% of the observed phenotypic variance (Fig. 4d, Table 1). The stickleback gene *Tnfrsf1b* is located within this QTL peak and is orthologous to the human gene *TNFRSF11A*, which is associated with susceptibility to Paget’s disease of bone in humans (Hughes *et al*. 2000; Makaram and Ralston 2021).

## Discussion

Although multiple previous studies have mapped genetic loci controlling the size, shape, and number of skeletal structures in stickleback, this is the first study to our knowledge to map differences in the internal microstructure of individual bones in the stickleback skeleton. Because high resolution µCT imaging and segmentation of individual bones in the skeleton is time- and labor-intensive, we were limited in the total number of animals we could score in the cross. Using phenotypic analysis of 102 F2 animals, we did not detect any significant QTLs for rib number, supraorbital groove or frontal bone sculpture prominence, armor plate bone volume fraction or mean bone thickness, or vertebral surface area or bone volume. One potential explanation is that a cross of this size only has sufficient power (>80%) to detect loci explaining 11% or more of the observed phenotypic variance in a trait (Purcell *et al*. 2003). We hypothesize, however, that many stickleback bone morphometry traits may be influenced by numerous loci each with small phenotypic effect rather than a few loci with large effects, consistent with findings from human GWAS studies of internal bone density (Ralston and Uitterlinden 2010; Richards *et al*. 2012; Morris *et al*. 2019; Zhu *et al*. 2021).

### Armor plate traits

Despite limited animal numbers, our genetic cross did detect significant QTLs for several armor plate and vertebral traits. Previous studies have shown that the number of armor plates in stickleback is controlled in large part by regulatory variation in the *Ectodysplasin* (*Eda*) gene on chromosome 4 (Colosimo *et al*. 2004; O’Brown *et al*. 2015). As expected, the absolute number of armor plates also mapped to the *Eda* region in our current cross. Several newly-measured aspects of the internal microstructure of armor plates also showed linkage to the *Eda* chromosome region, including surface area, total volume, bone volume, and mean bone separation (Fig. 3, Supplementary Fig. 3). One simple explanation for these results is that *Eda* plays a key role in plate development, and multiple different aspects of armor plate bone structure change because of pleiotropic consequences due to alterations affecting *Eda* itself. Selection for freshwater *Eda* alleles would then simultaneously reduce both the number of armor plates and the internal weight and mineral content of the bones, which may be beneficial in freshwater environments with less buoyant density and lower ion availability (Marchinko and Schluter 2007; Myhre and Klepaker 2009). Alternatively, there may be other linked genes near *Eda* that each contribute to different aspects of armor plate structure. A similar phenomenon has previously been reported for defensive spine formation in stickleback, where at least two different genes (*Msx2* and *Stc2a*) that are physically linked within 2 Mb of the *Eda* locus both contribute to genetic variation in length of dorsal and/or pelvic spines (Howes *et al*. 2017; Roberts Kingman *et al*. 2021a).

Interestingly, LOD scores along chromosome 4 show somewhat distinctive patterns for the different aspects of armor plate microstructure that we measured in the current study (Supplementary Fig. 3). In particular, while the peak marker for armor plate number and mean bone separation maps within the *Eda* gene itself, the peak marker for armor plate total volume and bone volume is ∼2.9 Mb downstream in the gene *Integral membrane protein 2A* (*Itm2a*). The peak LOD score for armor plate surface area falls between these two genes. *Itm2a* is involved in endochondral bone formation (Van Den Plas and Merregaert 2004), and either *Itm2a* or other physically linked genes in the chromosome 4 region may also contribute to one or more of the armor plate microstructure traits we investigated in this study. Indeed, in organisms as diverse as plants, butterflies, and stickleback, linked physical clusters of genes and regulatory elements (“supergenes”) are known to control multiple different aspects of complex flower, wing, and skeletal phenotypes (Schwander *et al*. 2014; Thompson and Jiggins 2014; Herbert *et al*. 2024). Our studies add to accumulating evidence that the *Eda*/*Itm2a* region of chromosome 4 is also a supergene region that controls multiple different aspects of armor plate development, defensive spine formation, and tooth and behavioral phenotypes (Erickson *et al*. 2018; Roberts Kingman *et al*. 2021a).

### Vertebral porosity

Three of the five vertebral bone morphometry traits showed modest but significant linkage signals, with total volume (TV), bone volume fraction (BV/TV), and mean bone thickness each mapping to chromosome regions with LOD scores just above the significance threshold. Two of these three vertebral traits also exhibited significant sexual dimorphism, with female fish displaying lower bone volume fraction (female mean: 0.6195, male mean: 0.6662, *p*_adj_ = 2.9 × 10^*-*13^ by Bonferroni-corrected Welch’s t-test) and lower mean bone thickness (female mean: −0.0029, male mean: 0.0028, *p*_adj_ = 0.0002 by Bonferroni-corrected Welch’s t-test), on average, than male fish. Both of these sexually dimorphic traits also mapped to chromosome 19—the stickleback sex chromosome (Peichel *et al*. 2004)—when sex was not included as a covariate in the QTL analysis (bone volume fraction peak at chr19:2,165,687, LOD: 6.7, PVE: 26.1%; mean bone thickness peak at chr19:2,344,544, LOD: 4.8, PVE: 19.5%). Inclusion of sex as a covariate eliminated these chromosome 19 signals, indicating that they reflect sexual dimorphism controlled by the sex chromosomes.

Bone volume fraction (BV/TV) of the 2nd vertebra—the proportion of the vertebra’s total volume that is mineralized bone rather than pores—is particularly interesting because of its relationship to osteoporotic changes in humans. Osteoporosis is characterized by increased bone porosity (i.e. lower bone volume fraction), and vertebral fractures are the second most common fracture type in osteoporosis patients (Compston *et al*. 2019). In humans, women who experience vertebral fractures in their 60s have ∼10% lower bone volume fraction (BV/TV) than age-matched women without fractures (Ito *et al*. 2005). In the stickleback we measured in this study, the mean bone volume fraction (BV/TV) was reduced by 3.3% from 0.664 in F2s with two marine BDGB alleles to 0.631 in F2s with two freshwater BOUL alleles at the peak marker (Table 1). Notably, stickleback genes with human orthologs related to bone development are located near the peak LOD scores for all three vertebral traits that pass genome-wide significance (Supplementary Fig. 4). These include human genes *CEBP, TNFRSF11A, ATP6AP1, SRC*, and *HSPG2* for total volume; *BBX, ACHE, PPP3CA, PANX3, FREM1, MEN1, SERPINH1, MMP14*, and *RBPJ* for bone volume fraction (BV/TV); and *CLEC3B, ECM1, ADAR, RXRB, LIMD1, SETDB1, HOXA11, CLEC5A, SNX10, NPPC, NR1I3, HEY1, PTK2*, and *TMEM107* for mean bone thickness of the 2nd vertebra.

While low bone volume fraction (i.e. high porosity) increases risk of bone fracture in humans (Ito *et al*. 2005), it may be beneficial in freshwater stickleback by helping to achieve neutral buoyancy in freshwater, which has approximately 2.5% lower density than ocean water (Myhre and Klepaker 2009), or by providing a growth advantage in environments with limited ion availability (Marchinko and Schluter 2007). Further study of stickleback genes within the bone volume fraction QTL may help identify genetic mechanisms that contribute to adaptive variation in skeletal density in different natural environments.

### Genetic basis of ivory vertebrae in stickleback

One unexpected finding in our study was the formation of unusually thickened “ivory vertebrae” in a fraction of F2 fish in the cross. This phenotype was not detected in either of the two F0 founders of the cross and occurred at relatively low incidence in subsequent generations (28 of 335 F2s). Unlike the relatively weak QTL for increased vertebral porosity, the QTL for presence of thickened ivory vertebrae is highly significant, with a LOD score of 12.9. At the peak marker, all but 4 of the 28 fish with ivory vertebrae are homozygous for the freshwater BOUL allele (3 are heterozygous, 1 is homozygous for the marine BDGB allele), suggesting that the freshwater BOUL allele acts as a risk allele in the cross.

Furthermore, the ivory vertebrae phenotype was observed in all four F2 families, with frequencies ranging from 7.8% to 9.0%, consistent with inheritance of a causative variant from the wild-caught BOUL founder through the shared F1 male and each of the four F1 females. The trait does not appear to cause significant genotype-dependent lethality, since genotype frequencies at the peak QTL marker for ivory vertebrae are consistent with the expected 1:2:1 Mendelian segregation ratio in the F2s (observed = 80:176:79, expected = 83.75:167.5:83.75, *p* = 0.65 by chi-square test). However, the trait also shows incomplete penetrance, with only 24 of 79 F2s (30%) homozygous for the freshwater BOUL allele exhibiting ivory vertebrae (Table 1), suggesting that while inheritance of the freshwater BOUL allele substantially increases susceptibility to ivory vertebra formation, additional genetic or environmental factors influence whether the phenotype is ultimately present..

The presentation of ivory vertebrae in this stickleback cross is reminiscent of Paget’s disease in humans, which is characterized by excessive bone resorption by osteoclasts that leads to excessive, disorganized bone formation (Makaram and Ralston 2021). The incomplete penetrance of ivory vertebrae in stickleback also parallels Paget’s disease in humans, where an interaction between genetic and environmental factors leads to disease presentation (Numan *et al*. 2019; Makaram and Ralston 2021; Banaganapalli *et al*. 2023). Genetic and genome-wide association studies of human Paget’s disease have identified more than a dozen different genes that increase risk of the disease (Makaram and Ralston 2021). One of these genes is *Tumor Necrosis Factor Receptor Superfamily Member 11A* (*TNFRSF11A*), which encodes Receptor Activator of Nuclear Factor Kappa B (RANK) (Hughes *et al*. 2000). Signaling through RANK is critical for osteoclastogenesis (Kong *et al*. 1999; Li *et al*. 2000; Boyle *et al*. 2003), and the human Paget’s disease mutations appear to be activating rather than inactivating mutations in the gene (Hughes *et al*. 2000; Alonso *et al*. 2021).

Interestingly, a stickleback homolog of this gene (*Tumor necrosis factor receptor superfamily member 1B, Tnfrsf1b*) is located within the highly significant chromosome 17 QTL for ivory vertebrae, less than 1 Mb away from the peak marker. The freshwater BOUL version of this gene shows 20 SNPs compared to the marine BDGB version based on sequence surveys of wild stickleback (Roberts Kingman *et al*. 2021b). Four of the BOUL SNPs are nonsynonymous variants that are predicted to alter two amino acids at positions otherwise conserved between marine stickleback and the distantly related medaka fish outgroup (Supplementary Fig. 5). As in human patients with Paget’s disease variants of *TNFRSF11A*, neither of these changes in *Tnfrsf1b* are predicted to be inactivating mutations (Ng and Henikoff 2003; Sim *et al*. 2012), but they could lead to more subtle changes in protein function. Alternatively, regulatory changes at the *Tnfrsf11b* locus, or coding or regulatory changes in other closely linked genes on chromosome 17, may contribute to the development of ivory vertebrae in stickleback.

Stickleback are often studied because of their usefulness for evolutionary and ecological questions. We show here that stickleback may also serve as valuable models for genetic studies of skeletal traits and disease that vary across a broad range of other animals, including humans.

## Supporting information

Supplementary File 1 - Genotypes and Linkage Map

Supplementary File 2 - Phenotypes

Supplementary File 3 - Coordinate Conversions

## Data availability

All linkage map, genotype, and phenotype data used for QTL mapping are provided as Supplementary Files accompanying this manuscript.

Supplementary File 1 contains the linkage map and genotype data for all F2 stickleback analyzed in this study. The first row lists linkage map marker IDs, named according to their physical position in the gasAcu1 genome assembly along with a unique identifier (e.g., the marker “chrI.1549902.SNP0006” corresponds to chromosome 1 at position 1,549,902 in gasAcu1, with SNP0006 as the unique marker ID). The second row provides the genetic position of each marker in centimorgans. All subsequent rows contain genotype information for individual F2 fish (IDs formatted as “DK184.013”). Genotypes are coded as “A” (homozygous for the freshwater BOUL allele), “B” (homozygous for the marine BDGB allele), and “H” (heterozygous, carrying one BOUL and one BDGB allele at that marker).

Supplementary File 2 contains ivory vertebrae status (0 indicates absence, 1 indicates presence), bone morphometric measurements, armor plate number, rib counts, and frontal bone phenotypes for all F2 stickleback. Both the original raw measurements (units: mm, mm^2^, and mm^3^) and the normalized residuals used for QTL mapping are provided for bone morphometric phenotypes. Residuals were calculated after multiple regression against standard length and sex. Raw and normalized values are included as separate columns (e.g. “armor-plate_total-volume” and “armorplate_total-volume.res” correspond to raw and residual values, respectively).

Supplementary File 3 provides a look-up table for converting select coordinates between the gasAcu1 and GAculeatus_UGA_version5 genome assemblies. Only coordinates cited specifically in the text or used in Table 1 and Supplementary Figures 3 and 4 are included. This file is provided because the linkage map marker names originally described in Wucherpfennig *et al*. 2022 and reproduced here in Supplementary File 1 were generated using gasAcu1 coordinates, whereas the Table and Figures have been updated to reflect the improved GAculeatus_UGA_version5 assembly.

## Author contributions

V.C.B. and D.M.K. designed the study. V.C.B. and D.L. performed phenotyping from µCT scans, and V.C.B. and J.I.W. performed phenotyping from standard X-rays. V.C.B. and J.I.W. conducted QTL mapping. V.C.B. and D.M.K. wrote the manuscript with input from all authors.

## Acknowledgments

We thank Brian Summers, Tim Howes, and Thomas Reimchen for earlier studies that laid the groundwork for the BOUL × BDGB cross; Kate Guenther for technical training on µCT; Amy Herbert for productive discussions on QTL mapping; and members of the Kingsley lab for valuable feedback and discussion.

## Author notes

Conflicts of interest: The authors declare that they have no conflicts of interest.

## Supplementary Figures

For Behrens *et al*. 2026, Genetic analysis of bone morphometry and ivory vertebrae in threespine stickleback

**Supplementary Figure 1:**
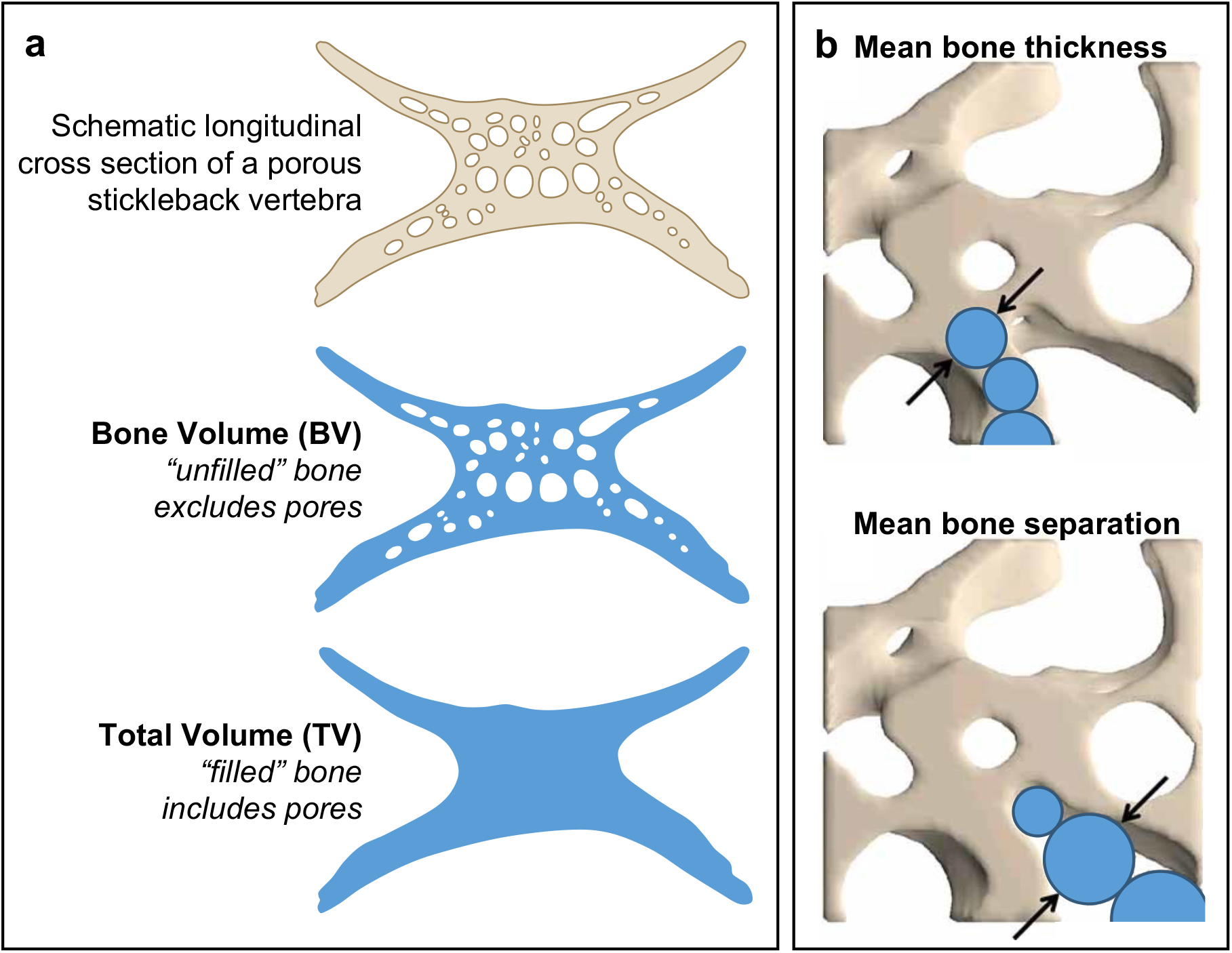
Schematic representation of μCT-derived bone morphometry measurements. **a**.Schematic 2D representation of the algorithm used to compute the bone volume (BV) of the unfilled bone-of-interest and the total volume (TV) of the filled bone-of-interest. Bone volume fraction (BV/TV) (not pictured) is the ratio of the unfilled bone volume to the filled total volume. Surface area (not pictured) is the external surface area of the unfilled bone. **b**.Mean bone thickness and mean bone separation are quantified by fitting spheres within the mineralized structure (for mean bone thickness) or within the porous space (for mean bone separation). The mean diameter of the fitted spheres is reported as mean bone thickness or separation. (Adapted from Fig. 6 of Bouxsein *et al*. 2010, originally courtesy of Andres Laib, Ph.D., Scanco Medical AG)

**Supplementary Figure 2:**
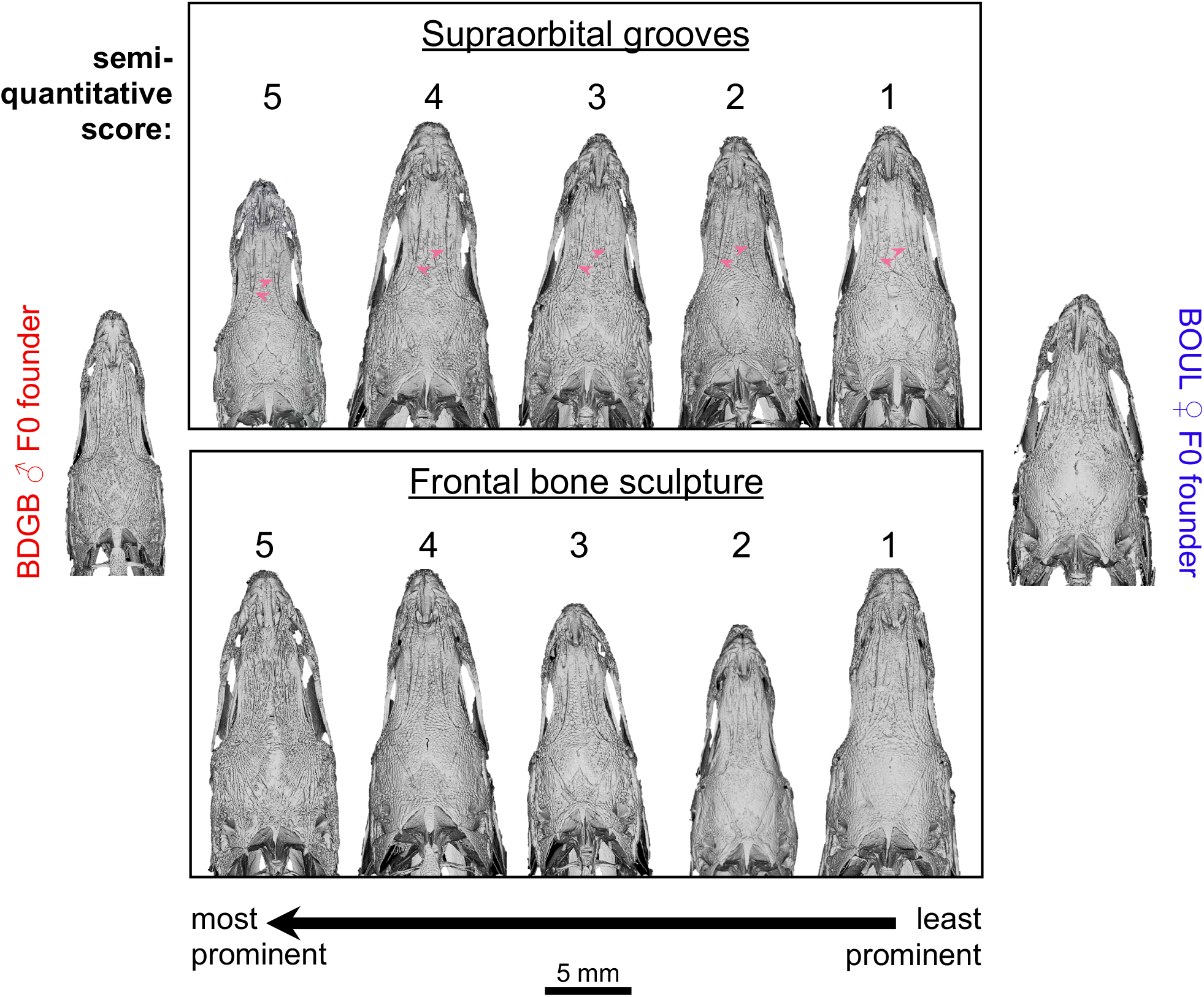
Semi-quantitative phenotype categories for supraorbital grooves and frontal bone sculpture. Dorsal view of representative F2 stickleback fish generated via μCT demonstrating the semi-quantitative scoring system for frontal bone traits. Supraorbital grooves (**top, arrowheads**) and frontal bone sculpture (**bottom**) were scored for QTL mapping on a scale from 1-5, with 5 indicating the greatest prominence. IDs for the F2s shown above are as follows, listed from left to right: Supraorbital grooves: DK194.007, DK185.125, DK185.003, DK185.131, DK185.005 Frontal bone sculpture: DK185.119, DK185.160, DK185.159, DK194.025, DK185.012

**Supplementary Figure 3:**
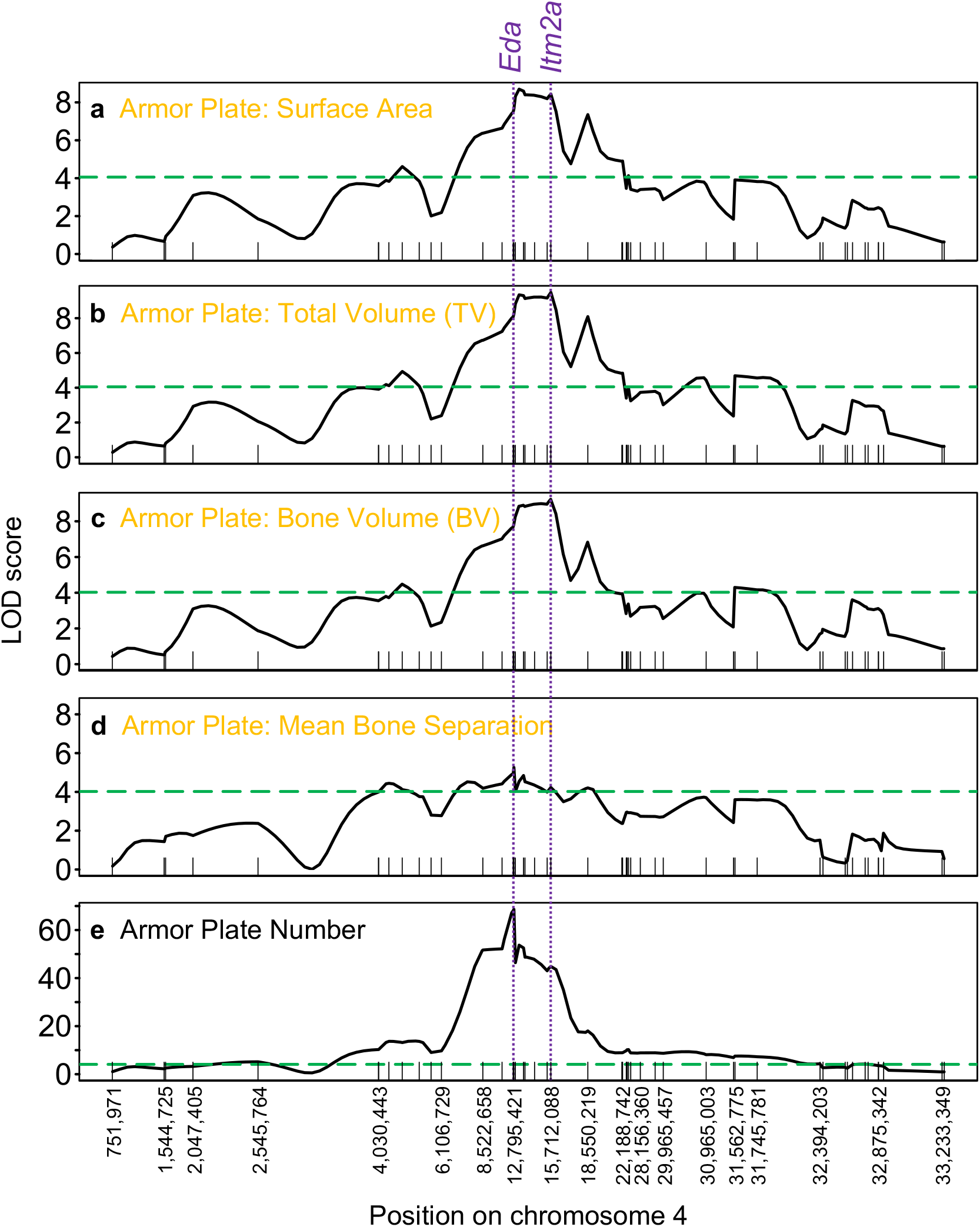
Candidate region genomic scans for armor plate QTLs. Single-chromosome QTL scan results for armor plate bone morphometry traits on chromosome 4 with significant QTLs. Dotted purple vertical lines spanning all plots indicate the position of the genes *Eda* and *Itm2a*. Note that the peak marker for total volume (**b**) and bone volume (**c**) is within *Itm2a*, but the peak marker for mean bone separation (**d**) and plate number (**e**) is near *Eda*. The peak LOD score for surface area (**a**) falls between *Eda* and *Itm2a*. Chromosome 4 coordinates (in the GAculeatus_UGA_version5 genome assembly) for select markers are shown on the y-axis, and plots were generated as described in the Fig. 3 legend.

**Supplementary Figure 4:**
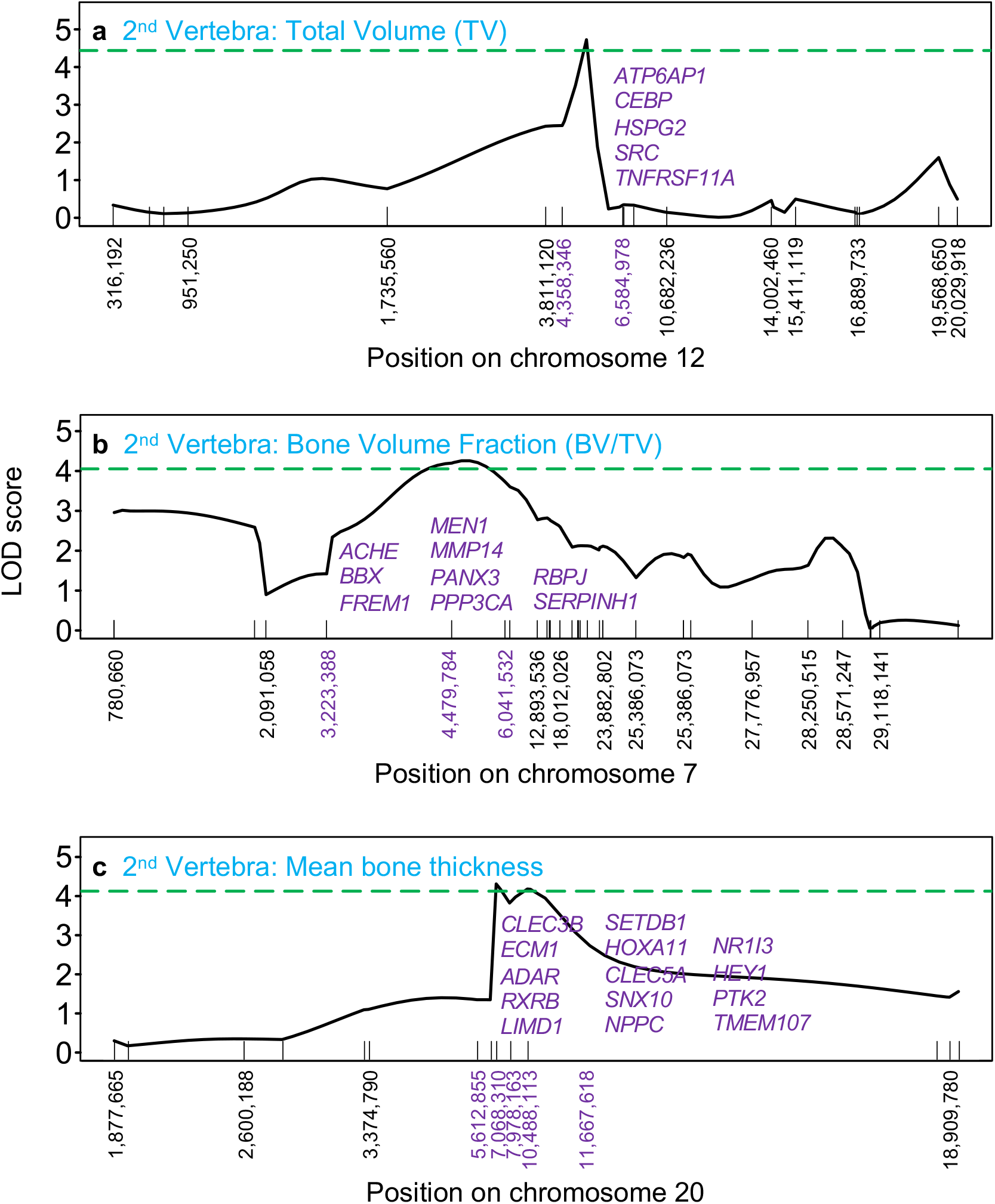
Candidate skeletal genes in 2nd vertebra QTL regions. Single-chromosome QTL scan results for 2nd vertebra traits with significant QTLs. Chromosome 12 (**a**), 7 (**b**), or 20 (**c**) coordinates (in the GAculeatus_UGA_version5 genome assembly) for select markers are listed on the y-axis. The coordinates in purple represent the candidate region used to identify candidate genes. Bone-related human orthologs for stickleback genes within the candidate region are shown in purple text on each plot. Plots were generated as described in the Fig. 3 legend.

**Supplementary Figure 5:**
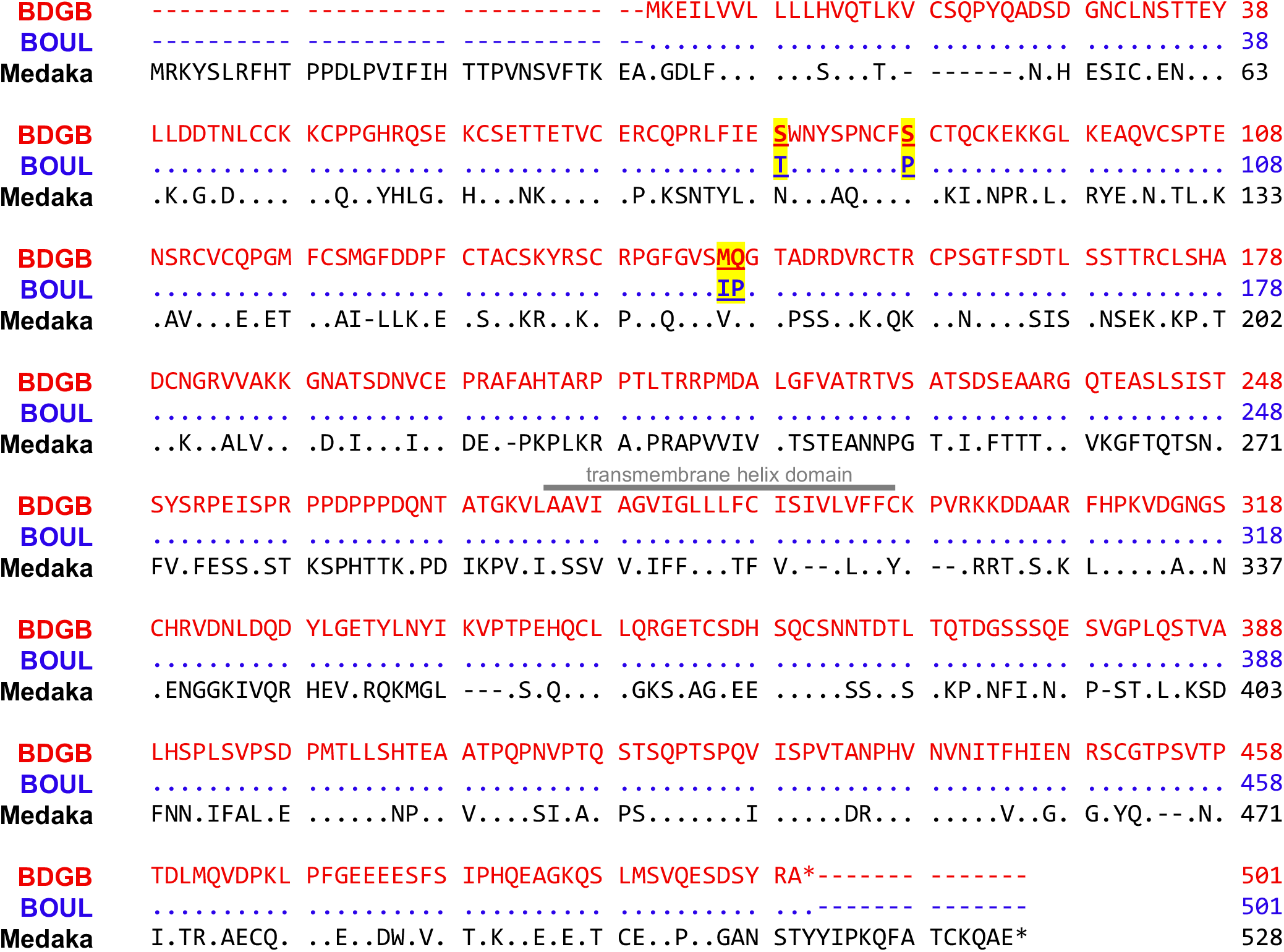
TNFRSF1B amino acid differences between BDGB and BOUL stickleback. Protein alignment of TNFRSF1B for the BDGB marine stickleback, BOUL freshwater stickleback, and outgroup Japanese medaka. Amino acid differences between BDGB and BOUL stickleback are underlined and highlighted in yellow. Two of the four differences are conserved to medaka. The transmembrane helix domain is indicated by a gray bar. The BDGB and BOUL stickleback protein sequences are based on ENSGACP00000029144.1 (transcript ID ENSGACT00000068214.1) and modified to incorporate amino acid changes resulting from population-specific SNPs identified from previously published DNA-sequencing data in Roberts Kingman *et al*. 2021b. The Japanese medaka protein sequence is ENSORLP00000005099.2 (transcript ID ENSORLT00000005100.2).

